# Steering From the Rear: Coordination of Central Pattern Generators Underlying Navigation by Ascending Interneurons

**DOI:** 10.1101/2024.06.17.598162

**Authors:** Julius Jonaitis, Karen L. Hibbard, Kaity McCafferty Layte, Atsuki Hiramoto, Albert Cardona, James W. Truman, Akinao Nose, Maarten F. Zwart, Stefan R. Pulver

## Abstract

Understanding how animals coordinate movements to achieve goals is a fundamental pursuit in neuroscience. Here we explore how neurons that reside in posterior lower-order regions of a locomotor system project to anterior higher-order regions to influence steering and navigation. We characterized the anatomy and functional role of a population of ascending interneurons in the ventral nerve cord of *Drosophila* larvae. Through electron microscopy reconstructions and light microscopy, we determined that the cholinergic 19f cells receive input primarily from premotor interneurons and synapse upon a diverse array of postsynaptic targets within the anterior segments including other 19f cells. Calcium imaging of 19f activity in isolated central nervous system (CNS) preparations in relation to motor neurons revealed that 19f neurons are recruited into most larval motor programmes. 19f activity lags behind motor neuron activity and as a population, the cells encode spatio-temporal patterns of locomotor activity in the larval CNS. Optogenetic manipulations of 19f cell activity in isolated CNS preparations revealed that they coordinate the activity of central pattern generators underlying exploratory headsweeps and forward locomotion in a context and location specific manner. In behaving animals, activating 19f cells suppressed exploratory headsweeps and slowed forward locomotion, while inhibition of 19f activity potentiated headsweeps, slowing forward movement. Inhibiting activity in 19f cells ultimately affected the ability of larvae to remain in the vicinity of an odor source during an olfactory navigation task. Overall, our findings provide insights into how ascending interneurons monitor motor activity and shape interactions amongst rhythm generators underlying complex navigational tasks.

## INTRODUCTION

Animal navigation requires coordination of different motor programmes over time. As animals navigate, the ratios of motor programmes controlling propulsion and direction are continuously adjusted to ensure efficient homing in on any goal. How animals navigate through environments has been extensively studied in both invertebrate ^1–4^ and vertebrate animals ^3,5–8^. However, the cellular mechanisms by which nervous systems coordinate and adjust motor programmes to enable homing in on navigational goals remain relatively poorly understood.

Directed movement requires coordination of rhythmic motor programmes underlying turning and propulsion. Central pattern generating (CPG) networks generate rhythmic motor patterns that underlie many locomotor programmes such as running, swimming, chewing and flying ^9–14^. Work in a variety of species has characterized the organization of CPG networks as well as how CPG activity is initiated, terminated, and modulated to enable effective movement ^9,15,16^. A traditional focus of CPG research has been on the mechanisms of rhythm and pattern generation ^17,18^ as well as intersegmental coordination ^19,20^ and modulation of speed ^21^. Recent work has begun to explore how descending and local interneuron populations can provide bilaterally asymmetric inputs to CPG networks to enable steering in both invertebrate ^22^ and vertebrate model organisms ^23^.

Interneurons that reside in higher order sensorimotor integration centers that send descending projections to motor circuits in either the spinal or ventral nerve cord are natural candidates for control of steering in locomotion^24–26^. However, interneurons that reside within motor centers and send ‘ascending’ projections to higher order centers could also play critical roles. A diverse array of bilateral ascending interneurons are present in both vertebrate and invertebrate locomotor systems and work in several systems have identified surprisingly complex functional roles for ascending interneurons, ranging from rhythm generation ^27^, to coordination of CPG activity ^28^, and even direct modulation of chemosensory inputs ^29,30^. Ascending neurons embedded within premotor networks and distributed across the anterior-posterior (A-P) axis of a locomotor network have the potential to shape flows of information about ongoing activity within a motor system. Bilaterally paired, ascending neurons are also inherently anatomically well suited to integrate information from motor centers and then in turn, influence bilaterally asymmetric activity in more anterior regions underlying steering in locomotion.

We used the *Drosophila* larval locomotor system to study how ascending interneurons interact with CPG networks to influence navigation. *Drosophila* larvae navigate towards and away from appetitive and aversive stimuli by generating a combination of bilaterally symmetrical propulsive peristaltic waves and bilaterally asymmetric head sweeps to sample their environment and change direction ^31–35^. A community led effort to generate a connectome based on electron microscopy (EM) data allows for identification of identified ascending neurons and comprehensive characterization of pre- and post-synaptic partners, while development of GAL4 driver lines allows for targeting of genetic tools for measuring and manipulating activity in specific cells and circuits ^32,36–40^. Previous work has identified cells and circuits critical for initiating or modulating forward and backward locomotion, stopping, turning, and integrating sensory information. For instance, multi-segmental ascending inhibitory neurons have been identified that play a role in promoting muscle relaxation ^41^ and locally ascending neurons have been shown to modulate speed or motor pattern formation ^42–44^. Recent work has also begun to identify the location and requirements for rhythm generating modules in this system. For instance, distinct CPG networks in posterior regions of the larval ventral nerve cord underlie forward propulsion^45^, while different sets of CPG networks underlie headsweeps. Intriguingly, in this system headsweeps are modulated by sensory input, but are driven fundamentally by rhythmic CPG networks ^46^.

Here we explored how a population of identified ascending neurons coordinate the activity of CPG networks underlying navigation in *Drosophila* larvae. We began by using neuronal circuit reconstructions from an EM volume to identify a population of ascending neurons with anatomical properties conducive to coordinating segmentally organized CPG networks. We then went on to explore the activity patterns and functional role of these neurons in locomotion and olfactory navigation. Overall, we found that these ascending neurons modulate the balance of forward propulsive movement with turning behaviour, which subsequently shapes how animals navigate in relation to odour sources.

## RESULTS

### Synaptic connections of 19f neurons

In *Drosophila* larvae, networks generating rhythmic patterns are distributed along the A-P axis of the ventral nerve cord (VNC), with regions toward the posterior primarily underlying forward locomotion and anterior regions predominantly coordinating turning movements ^32,36,47^. We reasoned that segmentally repeating, ascending interneurons would be well positioned to coordinate activity across CPG networks underlying larval navigation. To test this idea, we first used an electron microscopy volume of the 1^st^ instar *Drosophila* larval CNS to identify and reconstruct the anatomy and synaptic connectivity of a set of bilaterally paired interneurons termed ‘19f’ neurons (Figure 1). Paired 19f cells are present in all abdominal and thoracic segments and share several common features: 1) 19f cell bodies are positioned in a homologous position in each given hemi-segment and send axonal projections anteriorly; 2) 19f cell bodies and dendritic regions are located in the dorsal region of the VNC; 3) regardless of cell body location, all 19f cell projections terminate between the 2^nd^ and 3^rd^ thoracic segments; 4) no 19f axonal projections or dendritic processes cross the midline of the VNC; 5) 19f neurons do not contain large dense core vesicles, and 6) 19f cells do not receive direct inputs from primary sensory neurons (Figure 1). While tracing cell morphology, we identified pre- and post-synaptic sites on each 19f cell. Pre-synaptic sites were located primarily at the terminal regions of axons but were also present in clusters along axons within each hemi-segment (Figure 1A-C). Post-synaptic sites were located on neurites in the same segment as the cell body but were also present along axons and axon terminals (Figure 1A-C). Upon finding and reconstructing 19f cells across multiple segments, we found that at a population level, 19f neurons have an increasing number of output sites from posterior to anterior segments (Figure 1A, right panel). In contrast, at a population level, most post-synaptic sites found were in posterior regions of the CNS (Figure 1A, right panel).

**Figure 1:**
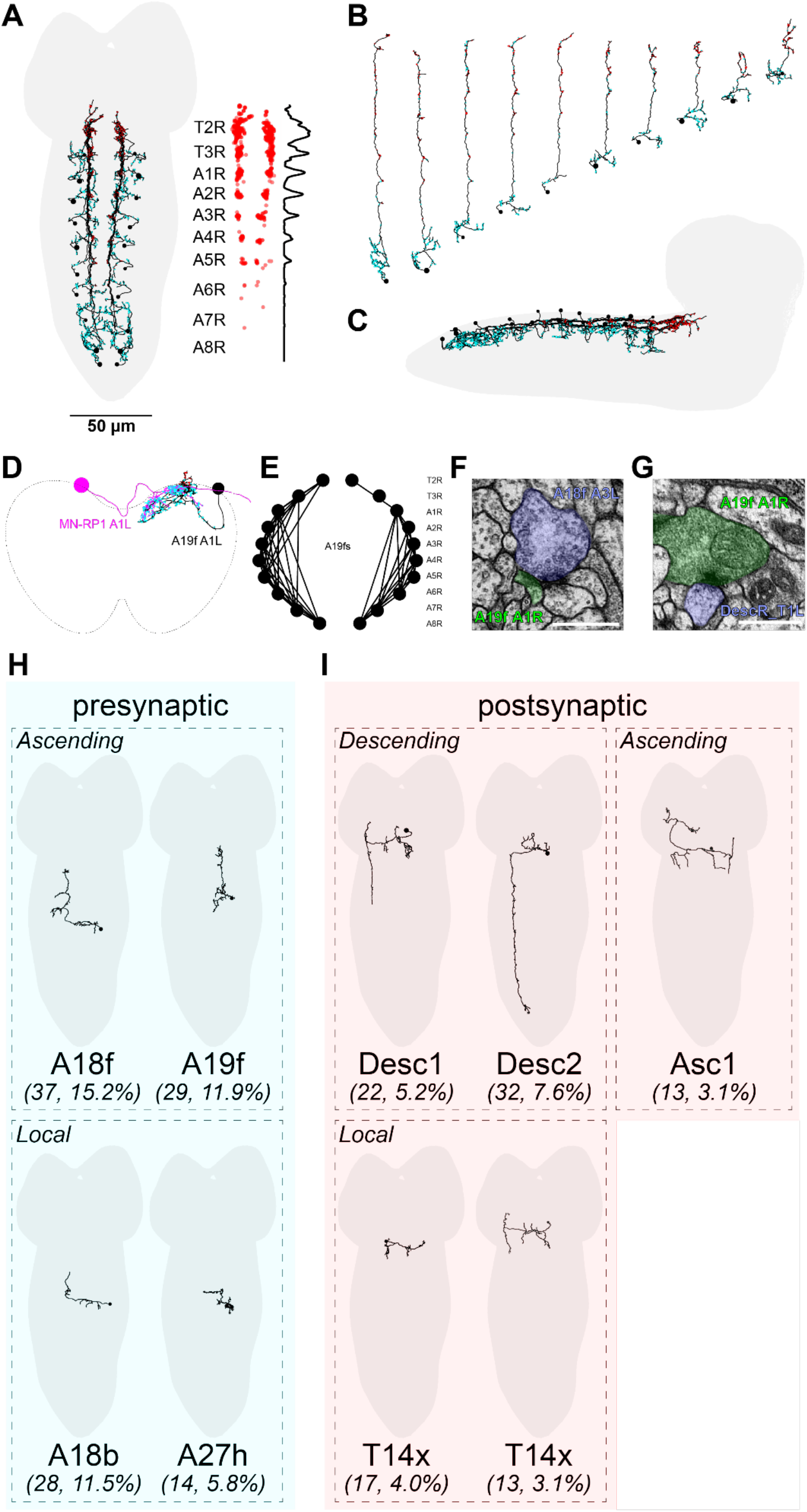
EM reconstruction of 19f anatomy in a 1^st^ instar larva **A**) Dorsal view of the reconstructed 19f cell population (black). Each segment contains one 19f cell that projects anteriorly. Red and cyan dots indicate pre- and post-synaptic sites respectively. **B**) Dorsal view of individual 19f cells on the left side of the ventral nerve cord (VNC). Each cell receives inputs primarily from the segment in which its cell body resides, and then projects to the first thoracic segment. Segmental identity of individual cells was determined using anatomical landmarks in EM volume (landmarks not shown, see text for details) **C**) Lateral view of the entire 19f cell population. Note that all neurites reside in the most dorsal regions of the VNC. Outline of the entire CNS is shown in light grey. Scale bar refers to all panels. **D)** Coronal view of 19f cells in relation to motor neuron in a single abdominal segment (A1L). Note lack of 19f processes in ventral regions. **E)** Schematic diagram of connectivity amongst 19f cells. Black circles represent cell bodies; labels indicate name and position of cell bodies in the VNC. **F)** Electron micrograph showing example of pre-synaptic sites to A18f_A3L (blue) to the A19f_A1R (green). **G)** Example showing post-synaptic partner DescR_T1L (blue) to the A19f_A1R (green) **H)** Showing examples of the most highly connected ascending and local presynaptic partners to A19f A1R cell (number and percentage of pre-synaptic connections to that partner). **I)** Examples of most highly connected descending, ascending and local post-synaptic partners to the 19f A1R cell body (number and percentage of post-synaptic connections to that partner).

After reconstructing pre- and post-synaptic partners of the 19f cell population, we observed that most highly connected pre-synaptic partners, including other 19f cells in different segments, reside primarily within the VNC (Figures 1E,H), and that 19f cells are strongly connected to one another axo-axonically (Figure 1E). The most highly connected pre-synaptic partners were other identified pre-motor interneurons^48^. This indicates strong interconnections among 19f cells and their involvement with other pre-motor interneurons. Another class of pre-synaptic partners were local pre-motor interneurons with axons that did not span across more than two abdominal segments (A18b and A27h) (Figure 1H). The 19f cell population’s post-synaptic partners also consisted of long-range descending interneurons as well as ascending interneurons with axonal projections that did not extend to the brain lobes, and several local pre-motor interneurons (Figure 1I). Finally, 19f cells also synapse onto neurons that project to brain regions. Overall, 19f neurons appear to receive inputs from a diverse array of pre-motor interneurons. In turn, post-synaptic partners of 19f are diverse and spread across multiple regions and cell types. The 19f cells are therefore well positioned to broadcast information about locomotor activity in the VNC to a variety of destinations and cell types in the larval CNS.

### GAL4 lines specific for identified 19f neurons

To examine the functional properties of 19f neurons, we created a 19f-Split-GAL4 line which targets 19f cells in 3rd instar larvae, with a small number of off-target ascending neurons in the thoracic region of the ventral nerve cord (Figure 2A, see Methods and Materials for details of line construction). This 19f-Split-GAL4 driver line drives expression in 12 19f cells, 6 on each side of the VNC (From A6 to A1), all 19f cells terminate between *subesophageal* ganglion segment 2 and 3. The off target thoracic cells terminate in the brain lobes, they also have processes projecting toward abdominal segment 2 and 3 (Figure 2A). Single cell labelling using a MultiColour FLPout approach revealed that 19f cells within 19f-Spit-GAL4 genotype closely match the morphologies of 19f cells found in EM reconstructions (Figure 2A, right side, see Methods and Materials).

**Figure 2.**
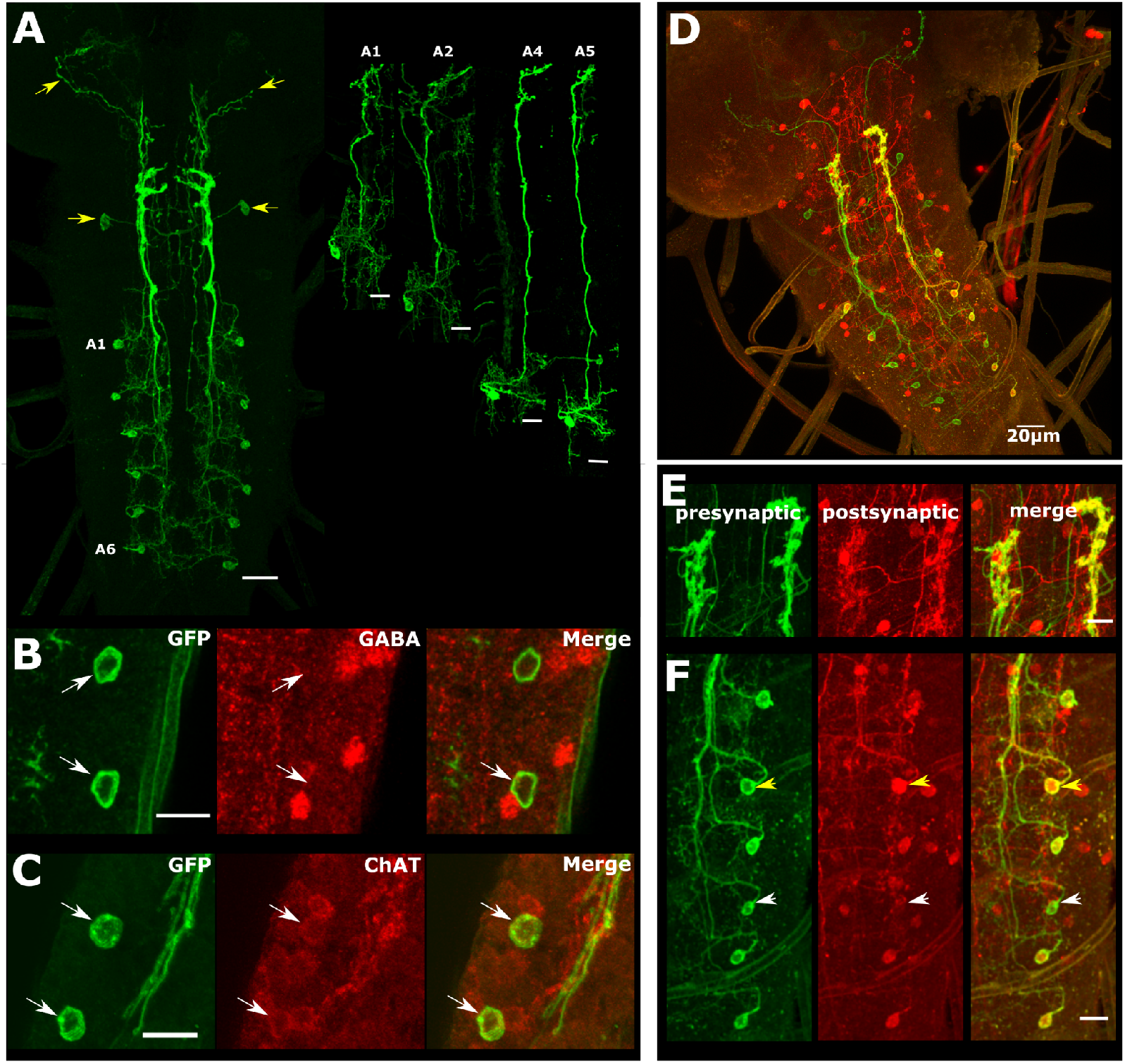
Expression pattern of 19f-split-GAL4, transmitter content and post-synaptic labelling of 19f cells using *trans*-Tango. **A)** Shows dissected 3^rd^ instar larval CNS with myrGFP expressed in 19f-Split-GAL4 driver line. 19f-Split-GAL4 drives expression in 12 19f cells, 6 on each side of the VNC, starting expression from abdominal segment 1 cell body up to abdominal segment 6 cell body (A1-A6). This driver line also has a few other ascending thoracic cells (yellow arrow) on both sides that are terminating in the brain lobes (yellow arrow, top left corner). Scale bar, 20µm. Right side panel shows single labelled 19f cells generated with Multi-Color-Flip-Out (MCFO) technique from different segments of the VNC (A1, A2, A4, A5). Scale bars, 10µm. **B)** 19f cells labelled with GFP (green) do not colocalize with cells stained for γ-aminobutyric acid (red) (GABA, n=3). **C)** 19f cells labelled with GFP co-localize with Choline Acetyltransferase staining (Red) (ChAT, n=3). Scale bars, 10µm. **D)** Expression of *trans*-Trango in 19f-split-GAL4 reveals multiple postsynaptic partners labelled with anti-HA antibody (red) to the 19f cell population labelled with anti-GFP antibody (yellow/green). Note that there are no motoneuron or sensory neuron as postsynaptic partners. Also there are no observed postsynaptic cells in the brain lobes. Scale bar, 20µm. **E)** Magnified view of the terminal regions of the 19f cell population also validates that these cells are downstream to one another (bottom right panel, merged). Scale bar, 10µm. **F)** Left panel, presynaptic, GFP expression in 5 19f cells (green). Middle panel showing labelled postsynaptic partners (red cell bodies and neurites). Third, top right panel showing merge (yellow) of the two channels. White arrowhead showing single 19f cell that did not colocalize. Scale bar, 10µm.

### Synaptic connections of 19f neurons in the 3^rd^ instar larva

Trans-synaptic labelling of post-synaptic partners of 19f cells using *trans*-Tango ^49^ (Figure 2D) confirmed that main anatomical features observed in EM reconstructions of 19f cells are also seen in 3^rd^ instar 19f cells. As in the EM dataset, most 19f cells are indeed connected to one another post-synaptically (Figure 2E,F), although there are some exceptions (note white arrowhead in Figure 2F). In addition, structural features of neurites visible in the EM dataset were also present in *trans*-Tango labelling (Supplemental Figure 1). To assay the transmitter content of 19f cells, we expressed GFP in 19f cells then performed antibody staining for either the neurotransmitter gamma-Aminobutyric acid (GABA) or the synthesis enzyme choline Acetyltransferase (ChAT). 19f cells did not show evidence of GABA immunoreactivity, but did show positive immunostaining for ChAT (Figure 2B,C, see Methods and Materials).

### 19f neurons are recruited into multiple fictive motor programs

Next, we wanted to gain insight into the functional role of 19f cells and examine endogenous activity patterns of these cells. To assess how 19f cells are recruited during fictive motor programs, we imaged Ca^2+^ signals in both 19f cells and aCC motoneurons by expressing genetically encoded calcium indicator GCaMP6f ^50^ in both populations (see Methods and Materials). Activity in the cell bodies of both populations as well as the terminal output regions of the 19f population can be measured separately (Figure 3A). During fictive locomotion, the cycle period of aCC activity peaks and the time delay from peaks in aCC to peaks in 19f activity was measured. This approach allowed us to determine whether individual 19fs were recruited and whether they preceded or followed motor activity within a given segment (Figure 3B). Both aCC motoneurons and 19f cells were active during fictive forward and backward waves (Figure 3D-G). 19f cells were also active in more anterior regions during bilateral asymmetries (Figure 3C), indicated with green or blue dots over asymmetric events (Figure 3F,G). Regions of Interest (ROI)s placed on the terminal regions of the 19f cells confirmed that 19f cells as a population were continuously active during waves observed in aCC motor neurons (top pink trace, TR, Figure 3D-G).

**Figure 3:**
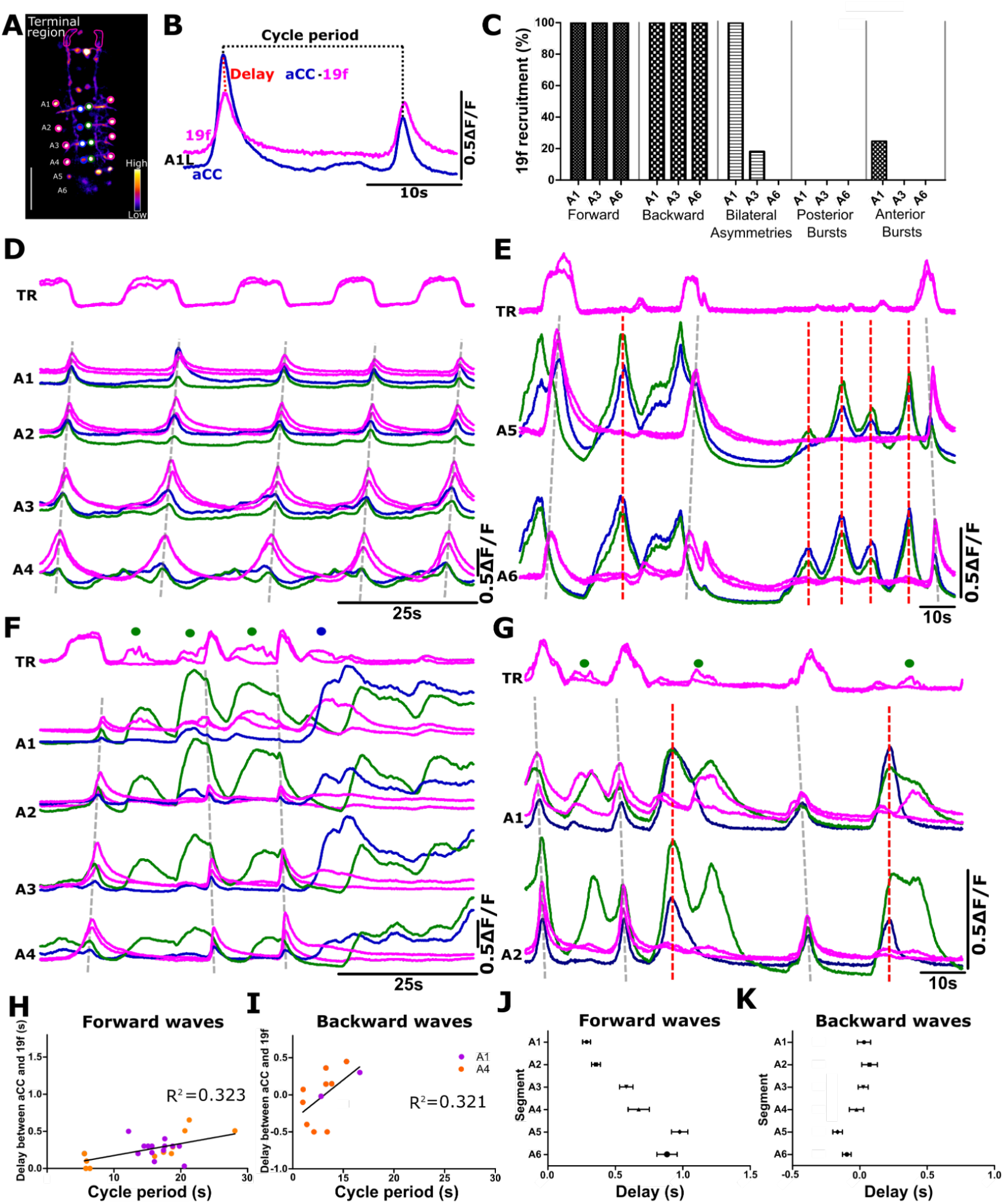
Simultaneous Ca^2+^ imaging of 19f cells and aCC motoneurons reveals that 19f cells are recruited into multiple motor programs. **A)** Single frame from an experiment in which Ca^2+^ activity is imaged in 19f cells and aCC motoneurons. ROI size and placement on motoneurons is shown on the right (green) and left (blue) hemi-segments. Same diameter ROIs were placed on 19f cells on both sides (pink circles) along with segment designations (to the left of image). Two more ROIs were placed on the terminal regions of the 19f cells (terminal region shaped, pink ROIs). Raw fluorescence is shown; differences in Ca^2+^ intensity changes are indicated by the colour lookup table bar at the bottom right. Scale bar, 100μm. **B)** Ca^2+^ signals of 19f cell and aCC motoneuron within A1 left side. Cycle period (peak of aCC activity to the next peak in the same segment) represented with dashed black line above trace. Delay between aCC peak time of activity and 19f cell activity within the same segment represented with dashed red line (peak of aCC activity to the peak of 19f cell activity). Scale bar (B), 50% change in fluorescence 0.5ΔF/F. **C)** 19f cells are active in all segments that were measured (A1, 3 and 6) during forward and backward waves and are active in most anterior abdominal segments during bilateral asymmetries. 19f cells are not active during synchronous posterior bursts and only few instances in 1/4 preparations where there was some activity in A1 19f cell during synchronous anterior burst. 19f cell recruitment in different fictive motor patterns is shown as percentage in different segments. Data collected from 8 different isolated CNS preparations. **D)** 19f cells are active in wave-like manner during forward waves propagating from most posterior segments towards more anterior (A4-A1). Terminal region (TR) showing overall activity of 19f cells as a whole population. **E)** Example trace where 19f cells were not active and motoneurons were producing synchronised posterior bursts of activity (red dashed lines) with few fictive forward waves (grey dashed lines). **F)** 19f cells are involved in fictive backward waves (in this trace first we see fictive forward wave followed by two backward waves) where activity starts in anterior regions and propagates towards most posterior. These cells are also active during bilaterally asymmetric events (denoted by green or blue dots depending on which side motoneurons were active), where only one side of the VNC is active. **G)** Comparison of 19f cell activity when motoneurons synchronously fire on both sides in the anterior regions and also produce fictive backward waves (grey dashed lines). **H)** 19f cells are active during fictive motor patterns with a short delay after aCC motoneurons fire during fictive forward waves, represented as duration of cycle period (peak of aCC burst to the next peak in the same segment) against the delay of 19f cell peak of activity in the same segment. Red line showing linear regression. **I)** Same as in **H**, but in fictive backwards waves. **J)** Showing delay in seconds between aCC motoneuron activity and the corresponding 19f cell activity from segments A1 to A6. Shortest delay between aCC peak of activity to 19f cell activity is in segment A1 and this delay increases across segments with the longest delay in most posterior segments. **K)** Same as **J,** but in fictive backward waves. Delay went into negative numbers which means that in few cases 19f cell activity preceded aCC motoneuron activity; n=5, error bars indicate SEM.

In all traces, Ca^2+^ signal data from left and right 19f cells was overlaid on top of one another to give an indication of whether 19f cells were active during bilaterally asymmetric events. 19f cells were recruited into most observable patterns of CPG activity; however, 19f cells were not recruited when aCC motoneurons produced synchronised bursts of activity in posterior regions that did not result in wave-like activity (note red dashed lines in Figure 3E). Most 19f cells were also silent when aCC motoneurons were recruited into active synchronous bursting in more anterior segments (red dashed lines, Figure 3G). In these experiments, we used a simple binary measure to gauge whether 19f cells were recruited in any given segment: 19f cells had to display a clear peak of activity with an amplitude that was at least 20% of the amplitude of aCC signals in the same segment. All 19f cells were recruited during all fictive forward and backward waves across all segments from A1 to A6 (Figure 3C). 19f cells in abdominal segment A1 were always active during bilateral asymmetries (Figure 3C). 19f cells in abdominal segment A3 were also recruited into bilateral asymmetries, but only during 18% of events (Figure 3C). In both A1 and A3, recruitment was always ipsilateral to corresponding aCC activity. 19f cells were not recruited during any synchronised posterior bursts, but 19f’s in A1 were recruited into 20% of synchronised anterior bursts (Figure 3C). Unlike synchronised posterior bursts of activity, synchronised anterior bursts occurred very rarely in preparations (activity pattern observed in 2/8 preparations, Figure 3E,G).

### 19f neurons are typically active shortly after motoneuron activation

To further investigate the timing of 19f cell activity relative to motoneurons, we measured cycle periods of aCC motoneuron activity in fictive forward and backward waves. Cycle periods were determined by measuring peak-to-peak times in aCC activity within a segment (Figure 3A). We measured peak 19f activity relative to aCC in A1 and A4 and found that 19f cells are usually active with a short delay following aCC activity across a range of cycle periods. There was considerable variability in delay times and no distinct scaling of delay relative to cycle period (Figure 3H). In fictive backwards waves, delay times were shorter and, in some cases, 19f cells fired simultaneously or even before motor neurons (Figure 3I). Next, we compared the delay between aCC and 19f activity across segments. 19f cells fired with a delay of 0.3 - 1s relative to aCC motoneurons in all fictive forward waves. Delays were not constant across segments, but rather increased from anterior to posterior regions (Figure 3J). In fictive backwards waves, 19f cell activity actually preceded aCC motoneuron activity in many posterior segments (A4-A6) and was marginally delayed in anterior segments (A1-A3) (Figure 3K).

### Optogenetic activation of 19f neurons modulates motoneuron activity

To explore how activating 19f neurons modulates CPG activity, we expressed CsChrimson in 19f cells and performed extracellular recordings in isolated CNS preparations exposed to red light pulses (see Methods and Materials). Loose cell-attached recordings from 19f cells subjected to 6 and 60-second pulses of 627 nm light revealed that both constant and high frequency light pulses evoke tonic spiking (Figure 4A-C). Pooled data for 6-second stimulations confirmed elevated spiking during and after the stimulation compared to quiescent control periods. (Figure 4D) (p < 0.05, p < 0.005, and p < 0.0001, respectively, n = 4). Once we had confirmed red light pulses would evoke elevated activity in 19f neurons, we performed suction electrode recordings from VNC nerve roots during ongoing CPG activity during 60 second red light pulses. Whether preparations were active or silent within a given region at light onset largely determined the type of response observed in that region. During moments of high activity within a region, activating 19f suppressed motor activity in that region; in contrast, during moments of silence within a region, activating 19f tended to elevate activity in that region (Figure 4E-H). To determine how activating 19f neurons affects posterior and anterior regions in isolation from one another, we transected CNS preparations then recorded from nerve roots in reduced preparations with brain lobes and thoracic regions intact, as well as preparations with only posterior abdominal segments present. Activating 19f in preparations without abdominal segments resulted in a decrease in the frequency of fictive head-sweeps. In contrast, activating 19f in preparations with only abdominal regions intact resulted in an increase in frequency of bursting (Figure 5). Taken together, these results suggested that the impact of activating 19f neurons is complex, being dependent on both the activity state of a segment as well as spatial location within the CNS.

**Figure 4:**
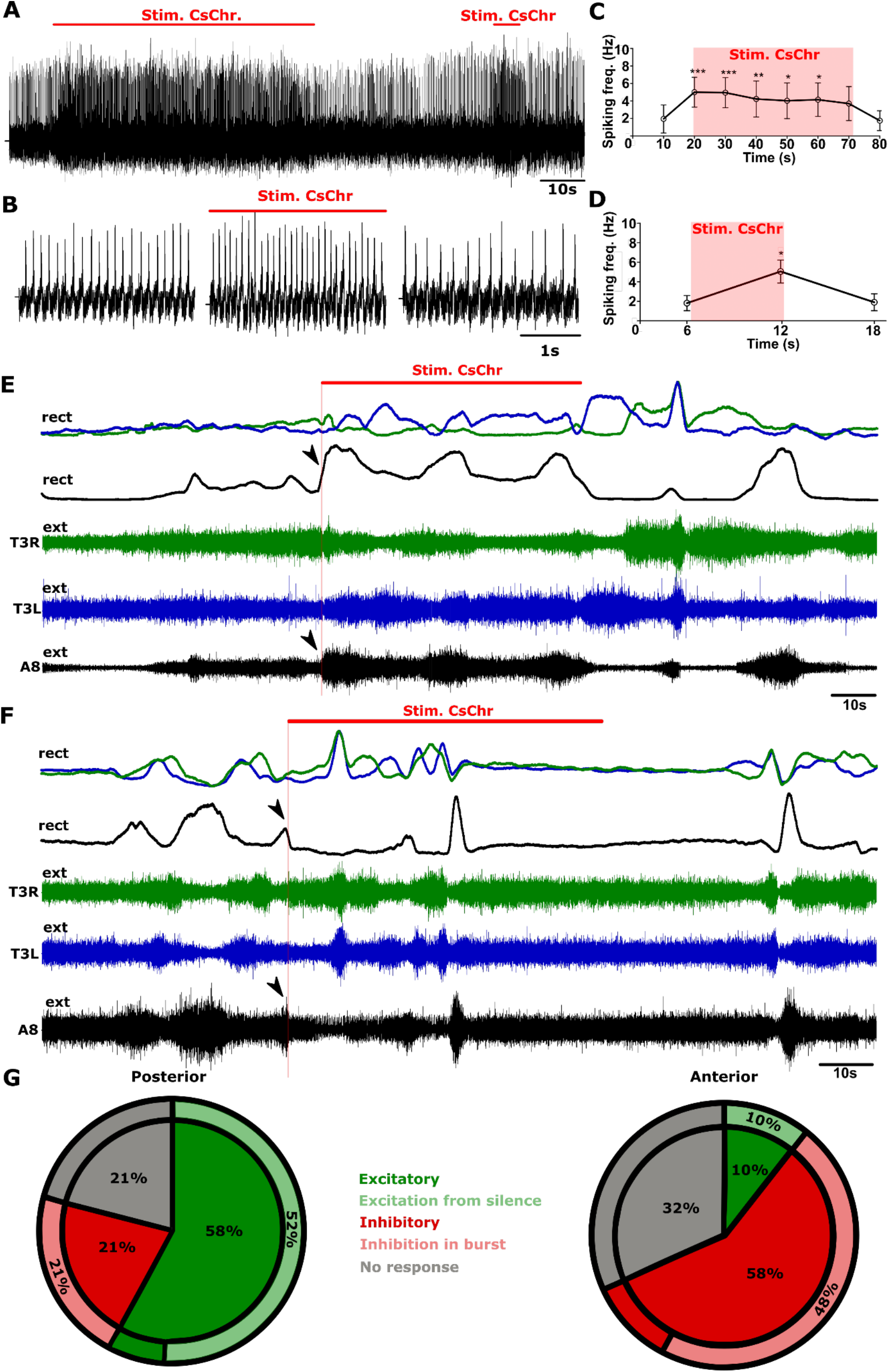
Activation of CsChrimson in 19f cells increases spiking frequency and also triggers state dependent changes in motor activity recorded in isolated larval CNS. **A)** Representative loose cell-attached recording from 19f cell with 6 and 60 second optogenetic stimulations (red lines above trace indicate time of light pulse). **B)** Enlarged view of periods before, during and after stimulation. **C)** Summarised spiking frequency data before, during and after the 60 second stimulation. 19f cell spiking frequency significantly increases during the red light stimulation. **D)** Pooled data for 6 s stimulations. Asterisks indicate significant differences between the before and during optogenetic stimulation (*P < 0.05, **P < 0.005, ***P < 0.0001; one-way repeated measures ANOVA with Bonferroni post-hoc test). n = 4. **E)** Extracellular suction electrode recordings from three nerve roots, two thoracic segments (green T3R and blue T3L) and one suction electrode recording from abdominal segment 8 (black A8/9). Extracellular nerve signals recorded from each electrode were rectified and smoothed by filtering with a moving average filter with a time constant of 0.9 s (rect). T3R and T3L overlaid to also observe bilateral asymmetries. Representative trace of motor neuron output while optogenetically stimulating 19f cell population for 60 seconds (red line indicates time of light pulse) showing excitatory response (black arrowhead) from the onset of the stimulation in posterior segment with inhibition in anterior segment T3R. **F)** Representative trace of motor neuron output while stimulating 19f cells showing inhibitory response in posterior segment also indicated by a black arrow, and excitatory response in anterior segment T3R. **G)** Pie charts representing percentage of different responses on the onset of 60 second stimulation in posterior regions (left chart) and anterior regions (right chart). n = 9, 18 trials.

**Figure 5:**
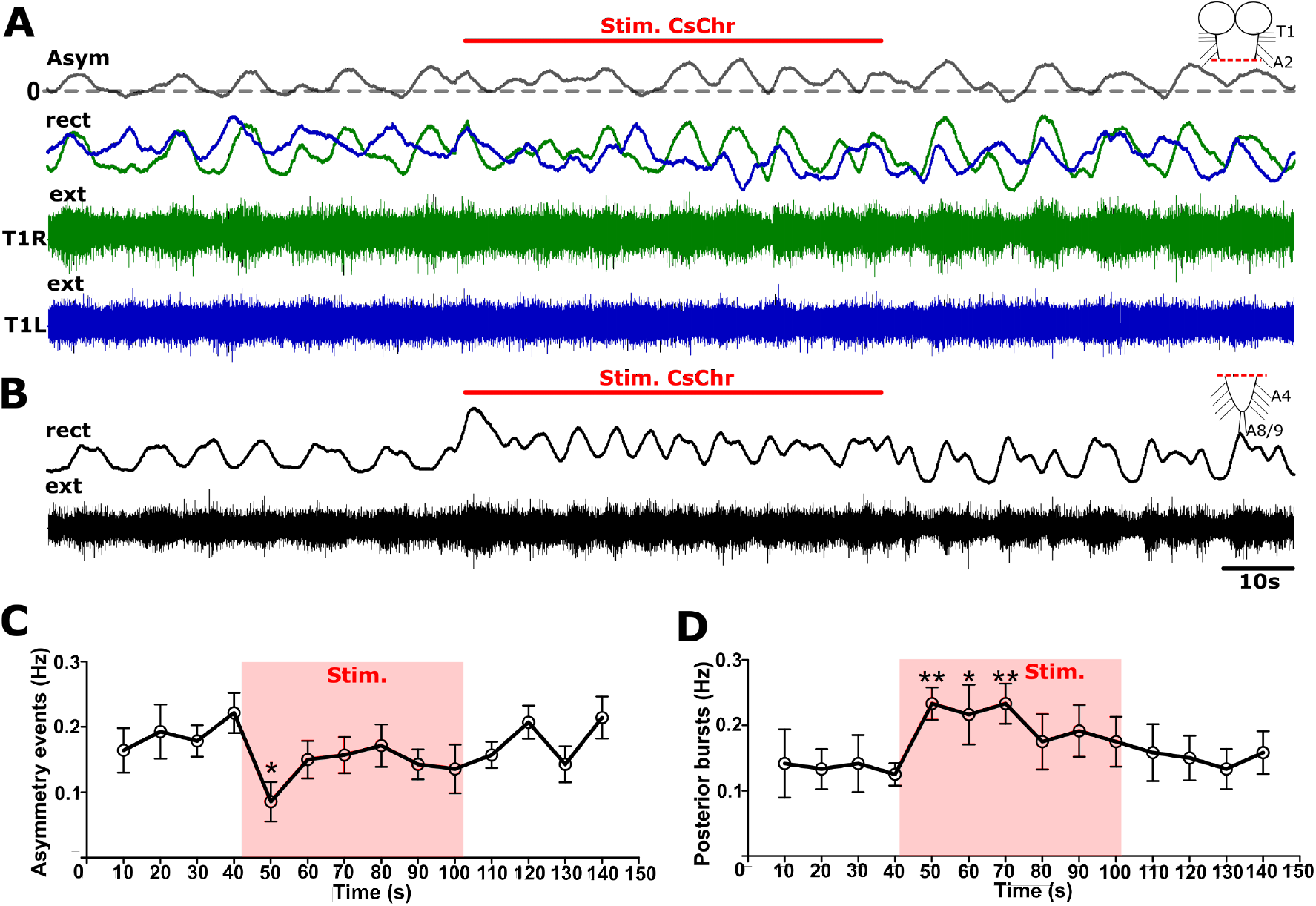
Optogenetic stimulation of 19f cells promotes rhythmic activity in isolated posterior segments and while supressing activity in isolated anterior regions **A)** Extracellular suction electrode recordings from two nerve roots, thoracic segments left and right sides (green T1R and blue T1L) in preparations where the posterior region is ablated and recorded from separately (black A8/9). **B)**. Extracellular nerve signals recorded from each electrode were rectified and smoothed by filtering with a moving average filter with a time constant of 0.9 s (rect). Rectified T1R and T1L outputs overlaid and bilateral asymmetries indicated by subtracting the signals from the two channels (Asym. grey line). Representative traces (**A,B**) shows motoneuron output while optogenetically stimulating 19f cells for 60 seconds in the two separated parts of the CNS. **C)** Summarised average frequency of bilateral asymmetries. **D)** Summarised average frequency of posterior end bursts. Asterisks indicate significant differences amongst frequencies at different time points (*P < 0.05, **P < 0.005, ***P < 0.0001; one-way repeated measures ANOVA with Bonferroni post-hoc test). Sample size, n = 7.

### Optogenetic suppression of 19f neurons does not alter motor activity

Next, we examined the effects of suppressing activity in 19f cells using the green and red light shifted optogenetic inhibitor GtACR1 ^51^. Simultaneous whole-cell patch recordings from 19f cells and paired suction-electrode recordings from nerve roots (A4L and A8/9) revealed that red light pulses effectively suppress spiking in these neurons (Figure 6A,B). We then conducted extracellular suction electrode recordings from two nerve roots in the thoracic segments (left and right sides) and from the abdominal segment 8 during red light pulses. Suppressing spiking activity in 19f neurons did not result in any clear activity dependent or location dependent trends. Overall, there were no clear effects on motor activity levels, nor on the frequency of fictive motor programs (Figure 6C,D).

**Figure 6:**
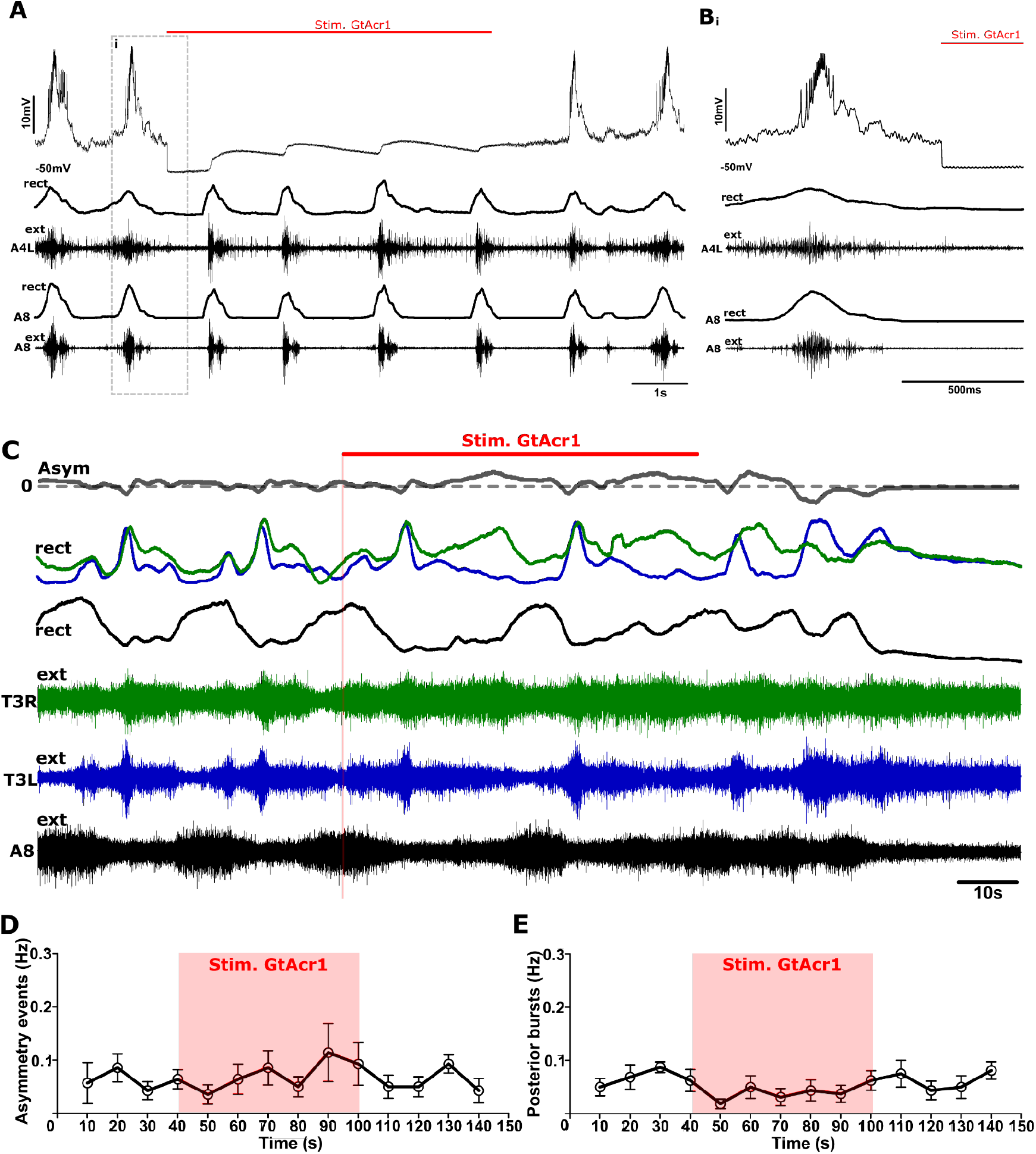
Activation of GtACR1 stops spiking in 19f neurons but does not have strong effects on network activity **A)** Representative trace of whole-cell patch recording from 19f cell recorded simultaneously with two suction electrode recordings from nerve roots (A4L and A8/9) (red line indicates time of light pulse). UAS-GtACR1 is expressed under the control of the 19f-Split-GAL4 driver line. **B)** Enlarged view of **Ai.** Representative trace. n = 2. **C)** Extracellular suction electrode recordings from two nerve roots, thoracic segments left and right sides (green T3R and blue T3L) and one from abdominal segment 8 (black A8/9) in intact preparations. Extracellular nerve signals recorded from each electrode were rectified and smoothed by filtering with a moving average filter with a time constant of 0.9 s (rect). Rectified T3R and T3L outputs overlaid and bilateral asymmetries indicated by subtracting the signals from the two channels (Asym. grey line). Representative traces shows motoneuron output while optogenetically inhibiting 19f cells for 60 seconds in the intact CNS. **D)** Summarised average frequency of bilateral asymmetries. **E)** Summarised average frequency of posterior end bursts. One-way repeated measures ANOVA with Bonferroni post-hoc test showed no significant differences between control and light pulse conditions. n = 7.

### Optogenetic activation of 19f neurons in freely behaving animals transiently suppresses locomotion

We next assessed whether manipulating the activity of 19f cells would impact the timing of locomotor behaviors (see Methods and Materials for details). First, we expressed CsChrimson in all 19f cells using the 19f-Split-GAL4 construct. Individual freely crawling experimental and control animals were stimulated with 60 second red light pulses. Ethograms of behaviour results were generated to visualize behavioural effects (Figure 7A). Light pulses triggered a halt in locomotion in experimental animals (Figure 7A, bottom ethogram) but had little or no effect on heterozygous controls (Figure 7A, top two ethograms). This halting behaviour was highly reproducible, but transient, lasting no longer than 6 seconds, regardless of the duration of stimulation (Figure 7B). During 60 second stimulations, forward wave frequency first dropped after onset of light pulse, but then started to return to baseline within 20 seconds (Figure 7B). Since the effects of stimulation were only seen during the beginning of the stimulation, only the first 30 seconds were considered for analysis. Forward wave frequency in experimental animals was significantly reduced (p<0.0001) compared to the controls and returned to control levels after the stimulation (Figure 7C). Forward wave durations during the first 30 seconds of the stimulation significantly (p<0.0001) increased compared to the controls (Figure 7D) and recovered to control levels after the light pulse. Headsweep frequency significantly decreased (p<0.001) during the first 30 seconds of stimulation compared to pre-stimulation controls (Figure 7E). Overall, these initial results showed that activation of the 19f cell population transiently interrupts larval crawling and affects both forward waves and headsweeps.

**Figure 7:**
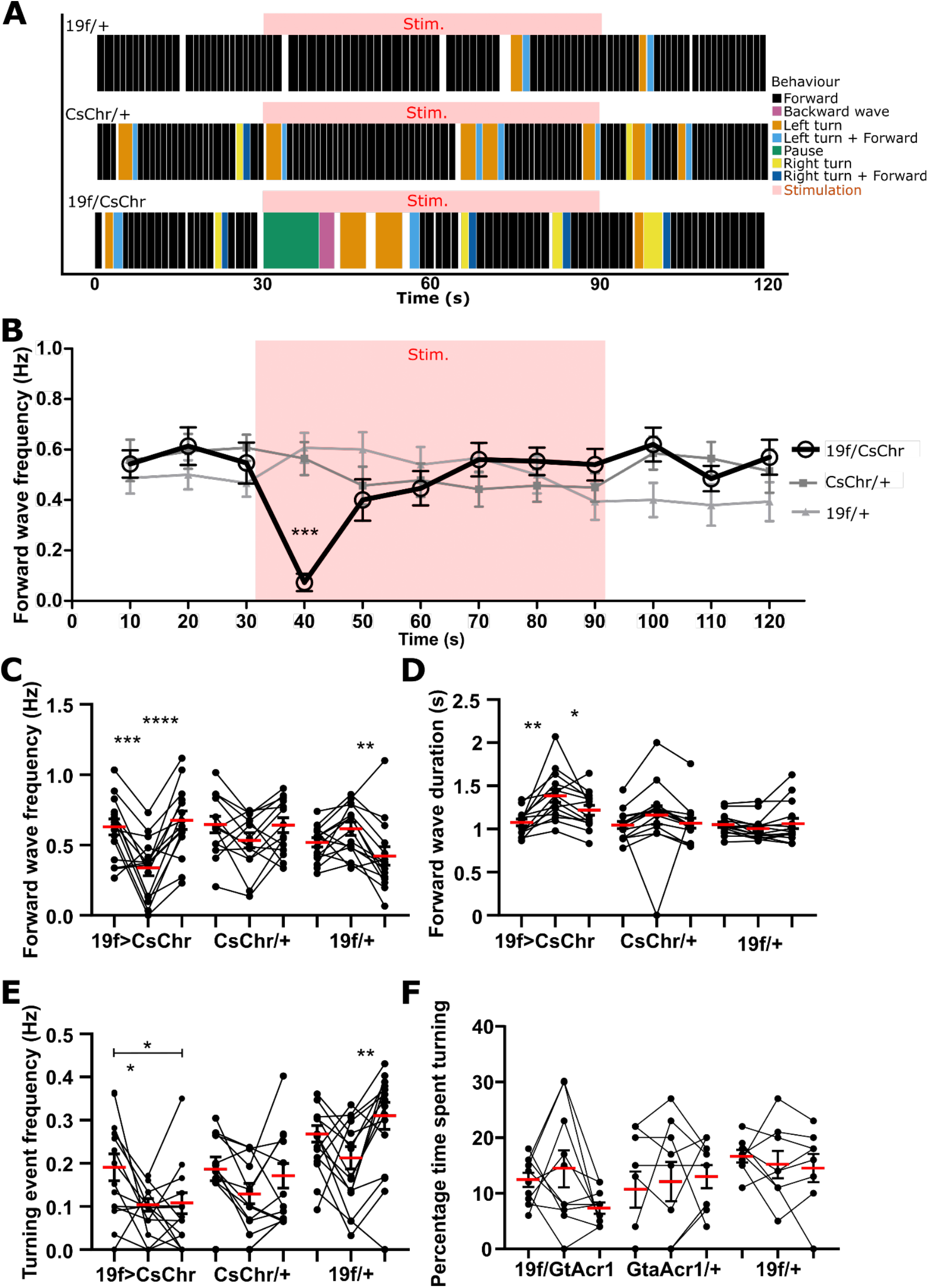
Optogenetic activation of 19f cells in freely crawling larvae leads to transient suppression of forward locomotion and turning behaviors **A)** Representative ethograms show different behaviours produced by freely crawling animals over time while optogenetically stimulating 19f cells (pink line represents onset and offset of red light). Different types of behaviours colour-coded in the figure legend on the right. The top two ethograms are the control animals (19f/+, CsChr/+) and bottom is experimental animal (19f/CsChr), where animal pauses for 5-6 seconds (indicated with black box) upon the onset of stimulation. **B)** Average forward wave frequencies of experimental animals (black line) significantly decreases upon the onset of stimulation (pink box represents start and stop of stimulation) compared to two other controls (grey lines). **C)** Summary of forward wave frequencies before, during and after the stimulation of experimental animals and two genetic controls. **D)** Summary of forward wave durations of experimental and control animals. **E)** Summary of turning event frequencies. **F)** Figure showing percentage of larvae spent turning. Asterisks indicate significant differences amongst groups (*P < 0.05, **P < 0.001, ***P < 0.0001; one-way repeated measures ANOVA with Bonferroni post-hoc test). Sample sizes were: n = 14 (19f/CsChr), n = 15 (CsChr/+), n = 14 (19f/+).

Our 19f-Split-GAL4 line expressed in a small number of cells that were not in the 19f cell population. In order to confirm that the observed activation effects are indeed being mediated by 19f cells (and not by off target cells in the expression pattern), we carried out Flp-Out optogenetic experiments in order to stochastically stimulate 19f cells alone (see Methods and Materials). CsChrimson was successfully expressed in different numbers of 19f cells ranging from 1 cell up to 6 in some preparations (Supplemental Figure 2A,B). In 48% of stimulations, either a short pause or wave interruption was present in animals with one or two 19f cells expressing CsChrimson (n = 5, 17 trials, Supplemental Figure 2C). The percentage of stimulations triggering pause or wave interruption increased to ∼75% in preparations where more than two 19f cells expressed CsChrimson (n = 6, 15 trials, Supplemental Figure 2C). A non-significant decrease in forward wave frequency was observed in both groups of animals (Figure 3D). A non-significant increase in duration of the first forward wave after stimulation onset was observed in the first group (1-2 19f cells, Supplemental Figure 2E) and a significant increase (p<0.05, and p<0.001) in first forward wave duration was seen in the second group (>2 19f cells, Supplemental Figure 2E). Overall, our Flp-Out optogenetics experiments produced phenotypes that were weaker but consistent with those seen in experiments using 19f-Split-GAL4.

### Optogenetic inhibition of 19f neurons in freely behaving animals promotes turning

We next inhibited 19f activity using the light gated anion channel GtACR1 to determine the impact of loss of 19f function on behaviour. In experimental animals, inhibiting activity in 19f cells induced an immediate slowing of forward wave propagation, and an increase in headsweep frequency (Figure 8A, bottom ethogram). Genetic controls showed no response to light pulses (Figure 8A, top ethograms). Forward wave frequency was significantly reduced for the first 20 seconds after the onset of light pulse (first ten seconds p < 0.0001 and p < 0.05 next ten seconds, Figure 8B) compared to both controls.

**Figure 8:**
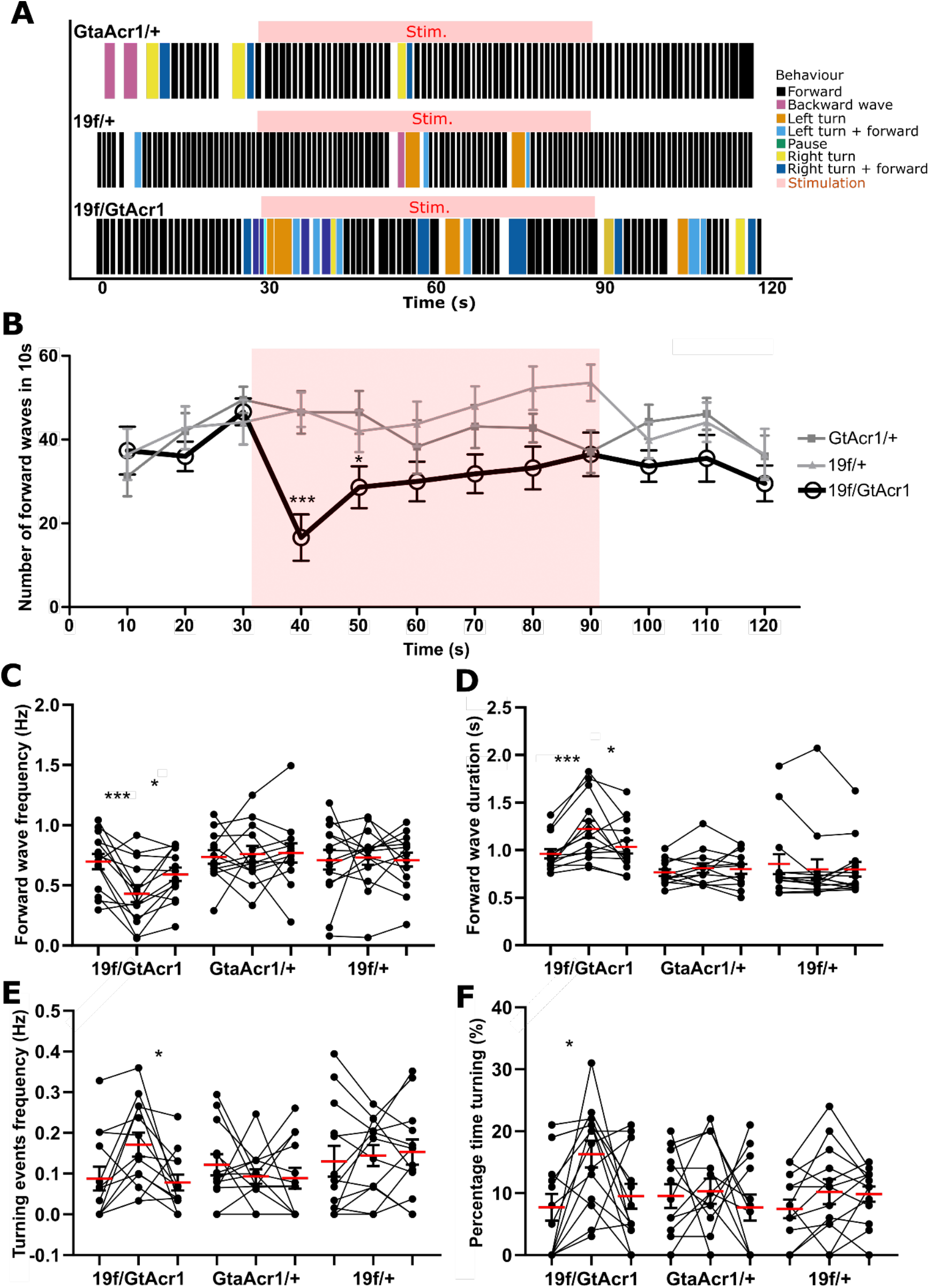
Optogenetic inhibition of 19f cells in freely crawling larvae suppresses forward locomotion but enhances turning **A)** Representative ethograms show different behaviours produced by freely crawling animals over time while optogenetically inhibiting 19f cells (pink line represents onset and offset of red light). Different types of behaviours colour-coded in the figure legend on the right. The top two ethograms are the control animals (GtAcr1/+, 19f/+) and bottom is experimental (19f/GtAcr1), where animal is doing more headsweeps during the inhibition of 19f cells. **B)** Average forward wave frequencies of experimental animals (black line) significantly decreases upon the onset of stimulation (pink box represents start and stop of inhibition) compared to two other controls (grey lines). **C)** Summary of forward wave frequencies before, during and after the red light of experimental animals and two controls. **D)** Summary of forward wave durations of experimentals against controls. **E)** Summary of turning event frequencies. **F)** Figure showing percentage of larvae spent turning. Asterisks indicate significant differences amongst groups (*P < 0.05, **P < 0.001, ***P < 0.0001; one-way repeated measures ANOVA with Bonferroni post-hoc test). Sample sizes were: n = 14 (19f/GtAcr1), n = 13 (GtAcr1/+), n = 14 (19f/+).

Experimental animals recovered in the second half of the light pulse and returned to control crawling after the light turned off. As in CsChrimson experiments, we only analyzed the first 30 seconds after the onset of the red light. Forward wave frequency was significantly decreased during the first 30 seconds of stimulation (p < 0.0001 and p < 0.001 compared to frequency after stimulation, Figure 8C). In addition, a significant increase in forward wave duration was observed during the first 30 seconds after red light onset (p < 0.0001 and p < 0.05 compared to before after the red light, Figure 8D). A non-significant increase in turning event frequency (i.e. headsweeps) was observed compared to pre-stimulation. However, a significant change in turning event frequency change was observed compared to post stimulation (p < 0.05, Figure 8E). Notably, compared to controls, experimental animals also spent significantly more time producing headsweeps and combinations of forward strides together with headsweeps (p < 0.05, Figure 8F).

### Constitutive suppression of 19f activity does not impede chemotaxis but reduces consummation of navigational goals

Our optogenetic manipulations suggested that 19f cells may influence both headsweep and forward wave generation in freely crawling larvae. To determine if 19f cells are influencing navigational programmes (which involve coordination of headsweeps and forward steps), we decided to assay how animals with inhibited 19f cells perform in an odor navigation assay. We inhibited activity in 19f cells by expressing the potassium channel Kir2.1 in the 19f-Split-GAL4 expression pattern, then examined how these animals navigate towards an attractive odorant compared to genetic controls (see Methods and Materials for details of assay). Most individuals in the experimental and control groups were able to successfully locate and reach the odor source (Figure 9A). A majority of animals, starting from ∼40 mm away from the odor came within a 15 mm radius of the odor source within 2 minutes (Figure 9B). Some animals, after finding the odor source, left and continued exploring (note blue lines in representation of animal distance from the center of odor source over time, Figure 9B). Mean ± SEM distance from center of odor source over time was plotted to visualize average navigational trajectories (Figure 9C). This did not reveal any drastic differences between experimental and control animals. The percentage of animals which reached the odor source at some point was similar across genotypes (Figure 9D). We then measured the total time that animals spent outside of the odor source (15 mm or more away from odor) which again showed no significant differences between the groups (Figure 9E). We noted that substantial numbers of experimental animals appeared to have reached the odor and then left (note blue lines in Figure 9B): 58% of experimental animals found, then escaped from the odor source, compared to 26 and 18% in genetic controls (Figure 9F). Finally, we counted the number of times animals left an odor source after finding it and found that experimental animals ‘escaped’ from the odor source significantly more times than one of the genetic controls (p < 0.05, Figure 9G).

**Figure 9:**
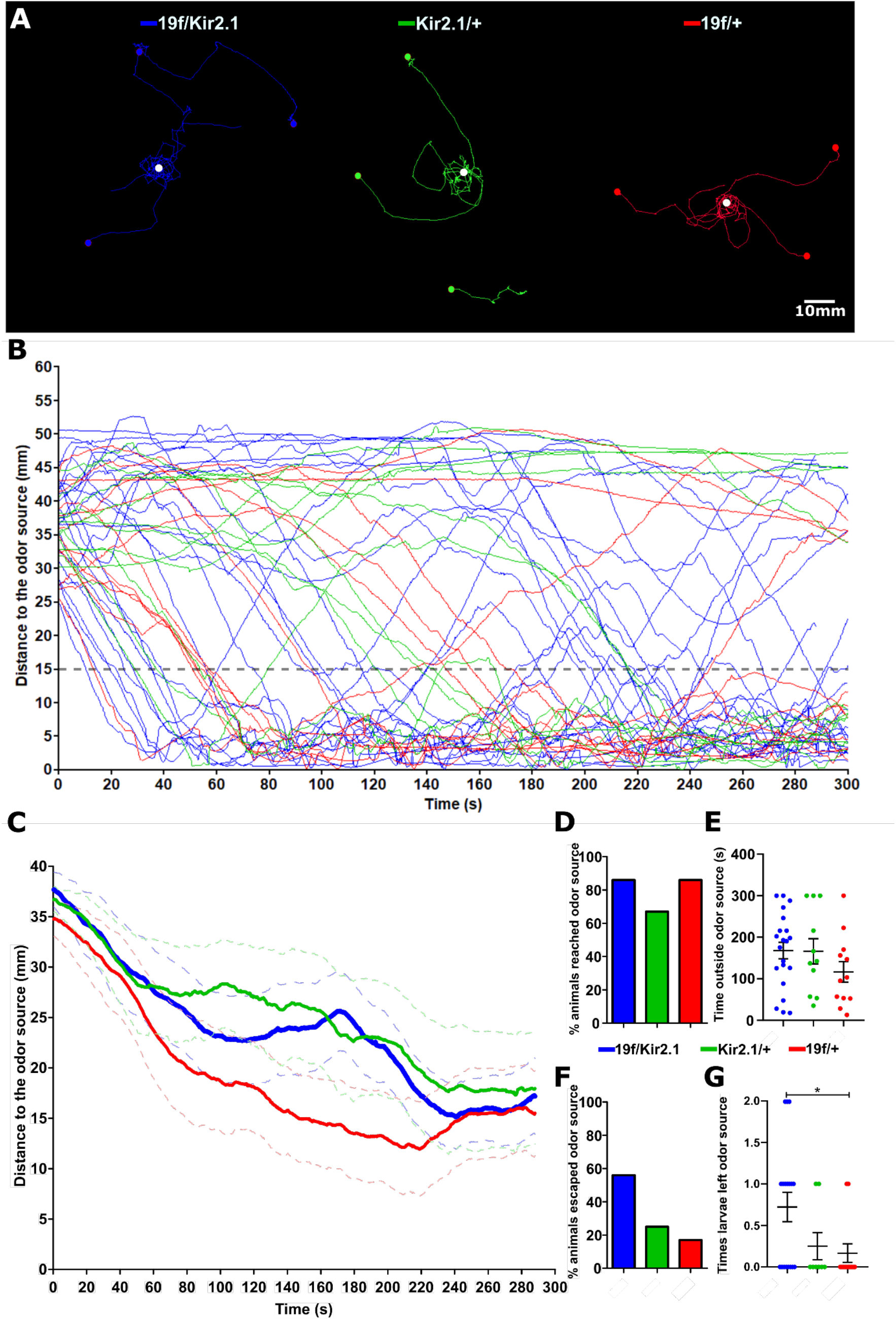
Expression of the potassium channel Kir2.1 in 19f cells disrupts the ability of larvae to remain near an attractive odor **A)** Representative images showing tracks of three larvae from their crawling starting point (starting point shown with coloured dots, different coloured tracks also represent different genotypes, legend above tracks). Animals were attracted by odour source (10μl drop) of ethyl butyrate (15mM) which was put into the center of clear glass cover (represented as white dot) of the behaviour arena so larvae can’t reach the odour source. **B)** Summary of all animals tracked which is represented as distance to the center of the odour source over time. Two controls were used in these experiments (Kir2.1/+ and 19f/+) and experimental animals (19f/Kir2.1). Grey dashed line shows 15mm distance to odor source. **C)** Averaged distance of larvae to the center of the odor source over time (dashed lines indicate ± SEM). **D)** Percentage of animals that reached within 15mm of the odor source. **E)** Total time animals spent outside of the 15mm region ±SEM. **F)** Representing percentage of animals that reached the odor source and then left the 15mm region. **G)** Number of times animals left the odor source outside of 15mm region ±SEM. Asterisks indicate significant differences amongst groups *P < 0.05; one-way ANOVA with Tukey’s multiple comparison test. Sample sizes were: n = 21 (19f/Kir2.1), n = 11 (GtACR1/+), n = 12 (19f/+).

**Figure 10:**
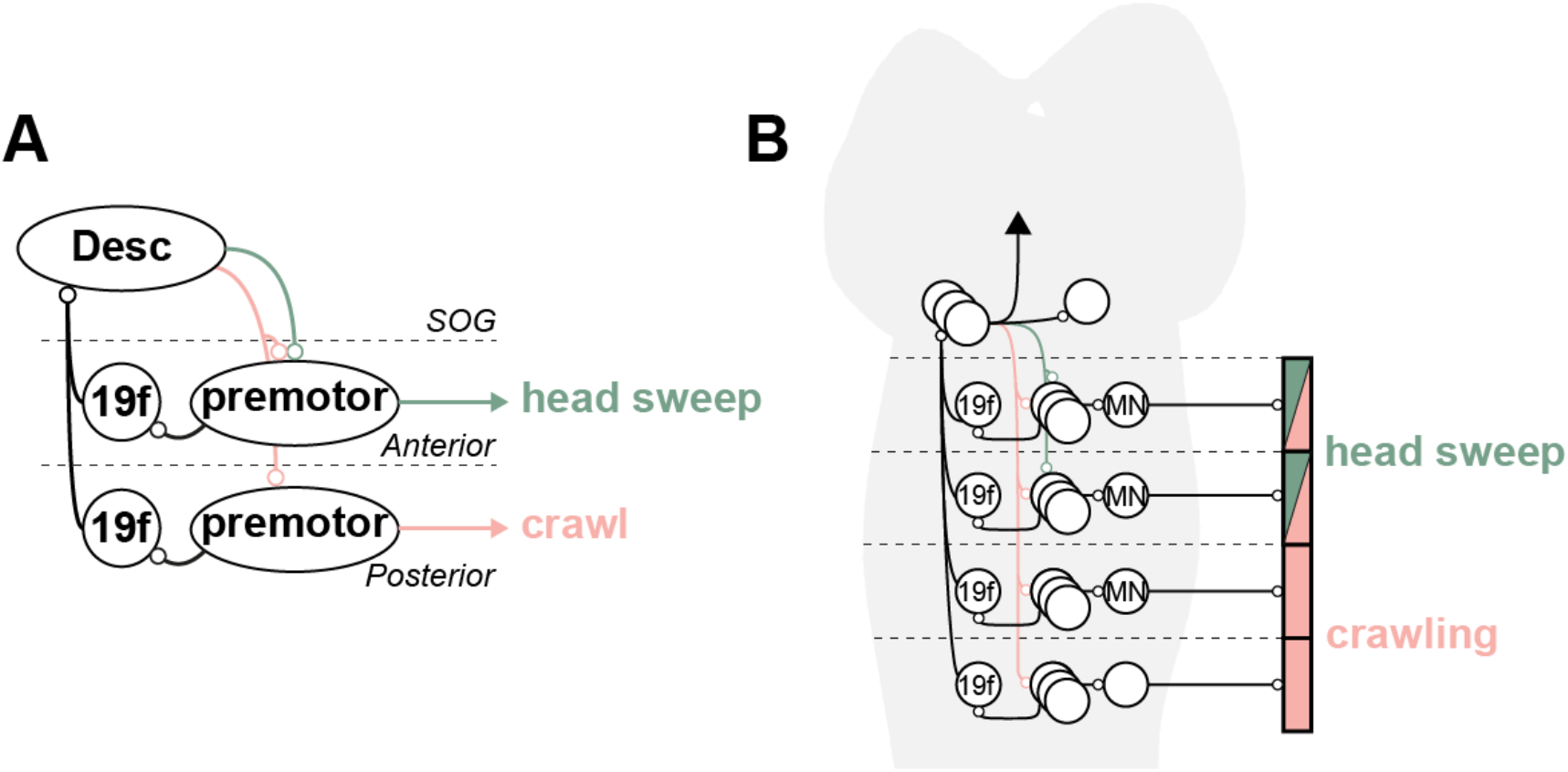
19f cells differentially modulate CPG activity underlying navigation in *Drosophila* larvae **A)** Diagram illustrating the neural circuitry involving 19f neurons (highlighted in red) and their connections to descending command neurons (Desc) and premotor circuits that orchestrate headsweep and crawl behaviors in *Drosophila* larvae. Activation of Anterior 19f neurons suppress headsweeps, while activation of posterior 19f neurons promote forward peristalsis. Modulating activity in the entire population has state-dependent effects that ultimately shift ratios of forward waves and turns. **B)** Topological representation of the 19f neurons within the central nervous system, arranged along the anterior-posterior axis. The organization reflects the division of labor in locomotor behaviors, with dedicated pathways for head sweeping (top) and crawling (bottom), as indicated by the red bars. This arrangement enables 19f neurons to monitor motor activity and modulate ratios of motor programs underlying navigation.

## DISCUSSION

Here we set out to determine the extent to which a population of ascending neurons embedded within a locomotor network shape goal-oriented navigation. We used a combination of anatomical, physiological, and behavioral analysis, to explore the function role of 19f ascending neurons in *Drosophil*a larvae. We show that these neurons influence ratios of motor programmes underlying navigation towards an attractive odorant - a task that is critical for survival in the wild.

The cholinergic 19f neurons synapse upon a large number of premotor interneurons, are presynaptic to a large diversity of cell types throughout the CNS and are highly interconnected with each other. Overall, these anatomical properties make the 19f cells well suited to gather spatio-temporal information about motor activity and then feed that information towards higher order brain centers in the CNS. 19f neurons appear to monitor motor activity, but notably, they do not have classical features of ‘efference copy’ neurons because they do not directly receive inputs from proprioreceptors and are therefore not immediately positioned to directly compare intended vs executed motor command ^52^.

Regardless, the terminal regions of 19f cells theoretically encode information about multiple motor programs at once. Previous work has suggested that these neurons could play a critical role in the formation of larval locomotor circuits early in development and this could reflect a role as motor activity monitors early in development ^53^. Neurons with similar types of morphology have been reported in larval ^27,41^ and adult *Drosophila* ^54,55^ as well as in vertebrate spinal cord ^56,57^, suggesting that these types of neurons could be conserved elements of locomotor networks. 19f neurons bear a resemblance to ascending histaminergic neurons (AHNs) in adult flies, which are also known for their integrative roles in sensory processing and motor control. Like 19f neurons, the AHNs feed information from motor centers to sensori-motor integration regions that modulate flight behavior ^58^. ‘Canon’ neurons in *Drosophila* larvae also show similarities to 19f, in that they project anteriorly from the segments in which they reside, are highly connected to one another and cholinergic. However, unlike 19f cells, canon cells show contralateral projections, and are recruited only into backward locomotor activity ^41^. Finally, results from whole central nervous system light sheet imaging revealed the presence of cells and regions that appeared to be monitoring different types of motor programs ^36^. Our work and the work of others suggest that in the larval locomotor system, different types of neurons can hold, and pass along information about the spatio-temporal patterns of motor programs. Exploring exactly how the larval locomotor systems uses information from these cell types to execute and adjust motor output is a potentially productive avenue for future research in this system.

Optogenetic activation of 19f cells triggered a complete, but transient pause in locomotion at the onset of stimulation; this was followed by lower frequency of headsweeps, and forward waves compared to controls. These experiments suggest that the main behavioral effect of activating 19f cells is to suppress locomotor activity. However, our work in isolated CNS preparations suggested that activating 19f cells could have either excitatory or inhibitory effects depending on network activity state – with inhibitory effects dominating during high activity states and vice versa. Locomotor rhythms in behaving animals are ∼10 times faster than fictive motor patterns in the isolated CNS, in part due to elevated levels of cholinergic excitation from sensory neurons ^32^. Given that VNC networks in behaving animals are in such a high excitability state, it makes sense that the effects of 19f would be generally inhibitory in intact animals. In our isolated CNS preparation, activation of 19f triggered different effects in different parts of the VNC. Activation suppressed activity in the rhythm generating modules that drive headsweeps and potentiated posterior rhythm generators that trigger forward waves. Our behavioral results in 19f activation experiments are consistent with the first observation but conflict with the second - headsweeps are indeed suppressed, but forward wave frequency is not potentiated. One possible explanation for the discrepancy is simply that the cycle period of the posterior rhythm generator is rate limited by wave activity – in this scenario, a potentiation of posterior rhythm generation could be masked by suppression of wave-like activity in other parts of the network.

How could activation of cholinergic, presumably excitatory, 19f cells generally suppress activity in the VNC? Our EM reconstruction shows that multiple GABAergic interneurons are postsynaptic to 19f; activation of these partners in concert could potentially have widely inhibitory effects. Some inhibitory partners are also known to be directly premotor. Inhibitory A26f neurons for example are directly pre-motor to a subset of motor neurons that selectively innervate transverse muscles, a functionally distinct subset of body wall muscles ^59^. Identified GABAergic neurons downstream of 19f cells in thoracic and subesophageal regions would be ideal entry points for further explorations aimed at revealing core circuits for generating headsweeps.

Hyperpolarizing 19f neurons with GtACR1 led to increased time spent in headsweeps. One possible explanation for this result is that the loss of 19f cells within the network leads to maladaptive timing overlaps in headsweeps and forward waves. A working model is that 19f cells are recruited during forward waves and their activity suppresses the generation of headsweeps. When this suppression is removed during GtACR1-mediated silencing, head sweeps are generated prematurely in the step cycle, leading to ‘over turning’ and navigational instability. The inhibition of forward wave speed during optogenetic inhibition is at first somewhat confusing, given that the same phenotype was present when 19f cells were activated. This combined phenotype is consistent, however, with 19f cells playing a role as ‘wave monitors’ within the larval CNS. If 19f cells are monitoring motor wave activity, then it is conceivable that any disruption in their activity could disrupt the network’s ability to be aware of what multiple segments are doing at any given time, resulting in slower, more irregular wave propagation across segments.

Our optogenetic experiments supported the idea that 19f cells function to coordinate rhythmic locomotor patterns in different regions of the VNC. In our final set of experiments, we assayed odor navigation in controls, and in animals with constitutively inhibited 19f cells. Successful *Drosophila* larval chemotaxis requires coordinated transitions between different types of behaviours (forward crawling, stopping, lateral headsweeps and asymmetrical contractions followed by straightening)^60,61^. We reasoned that animals with inhibited 19f cells may have difficulties with chemotaxis, given our headsweep phenotypes in GtACR1 experiments. We found that experimental animals, just like genetic controls, were still capable of locating and reaching odor sources. But experimental animals, after reaching odors, left and in some cases re-entered the odor area again significantly more times than control animals. These results suggest that unlike recently identified interneurons (e.g. ‘PDM’ neurons; ^62^), 19f neurons are not directly involved in coupling the olfactory system to motor actions during initial approaches to odor sources. 19f cells appear to play a more nuanced role in later stages of olfactory navigation. Our findings suggest that these neurons are playing a role in balancing the need for forward propulsion with headsweeps to achieve navigational precision. This integration enables the larvae to adapt their movement strategies in real time—suppressing or enhancing different ratios of motor programs as needed to optimize interaction with their environment. The ability of 19f neurons to control such diverse behavioral outputs positions them as key components in the sensory-motor pathways that govern adaptive behavior in response to dynamic sensory landscapes. This raises the interesting possibility that neurons and neuronal populations with similar anatomical and functional motifs could be playing similar roles in other species.

Overall, our findings contribute to a broader understanding of the neural circuitry underlying locomotion and motor program flexibility during navigation, showcasing the role of ascending interneurons in these processes. Unravelling the roles of these types of neurons in *Drosophila* provides a potential roadmap for discovering conserved principles governing how circuits embedded deep within locomotor networks can steer from the rear and influence how animals navigate through sensory landscapes.

---

**Supplemental Figure 1:**
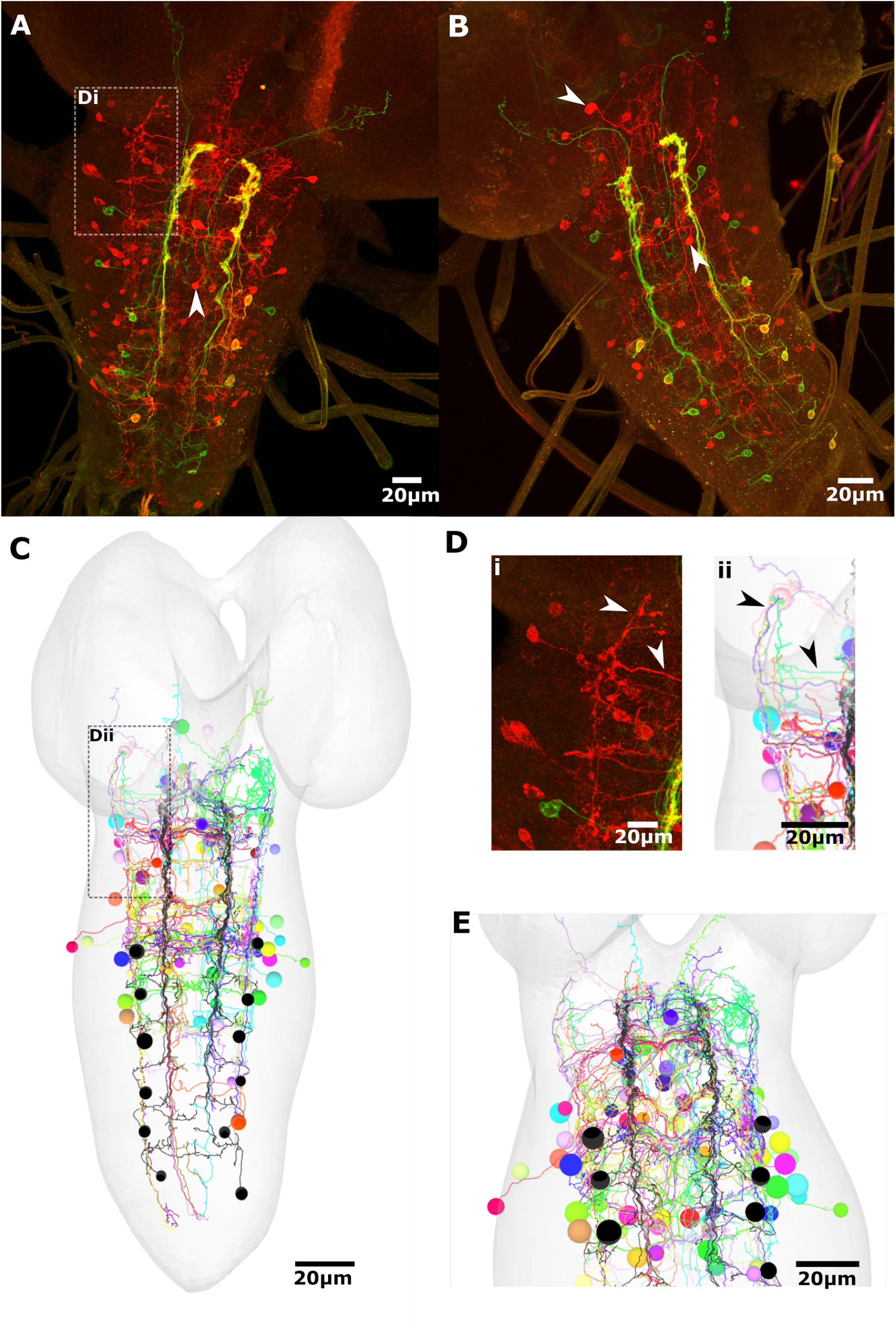
Comparison of *trans*-Tango postsynaptic labelling of 19f cells against EM reconstructions of 19f cell postsynaptic partners. **A,B)** Two different preparations (as in figure 2) expressed *trans*-Tango in 19f-split-Gal4 and labeled postsynaptic signal with anti-HA antibody (red) and 19f cell population labeled with anti-GFP antibody (yellow/green). Similar morphological features can be observed in both preparations, including already mentioned absence of motor neuron partners. **C)** EM reconstructions of 19f cells from A1 to A6 and their post-synaptic partners connected by at least 3 synapses also resemble some of the main morphological features observed in *trans*-Tango staining. **D)** Magnified views of descending, ascending processes observed in both *trans*-Tango staining (**i**) and corresponding magnified view of EM reconstructions (**ii**) main resembling morphologies denoted with arrows. **E)** Enlarged view of thoracic, anterior abdominal segment reconstructions of post-synaptic partners to 19f cell population. All scale bars, 20μm.

**Supplemental Figure 2:**
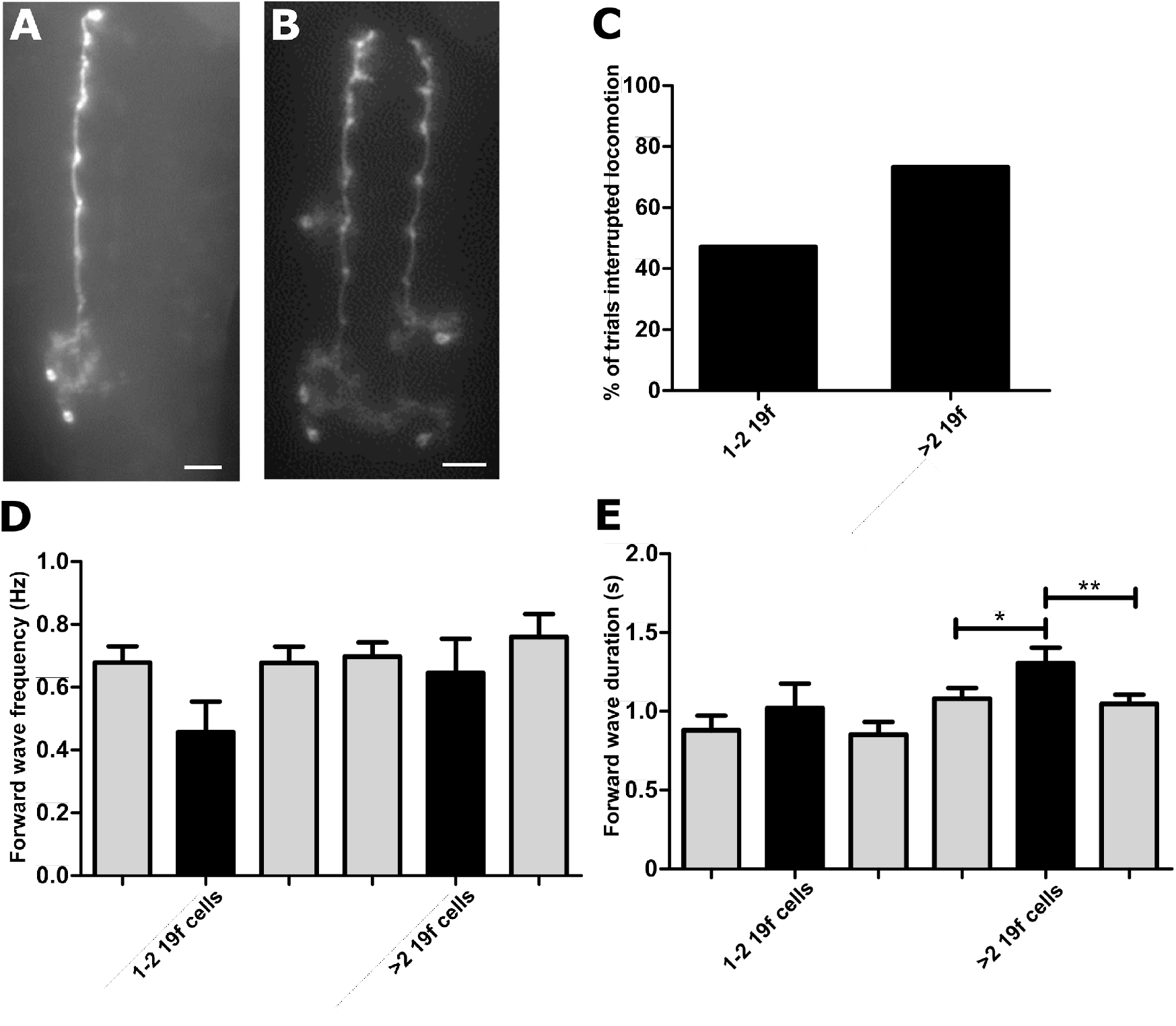
Activation of small numbers of 19f cells using FLP-out optogenetics produces phenotypes consistent with activation of the full 19f-GAL4 expression pattern **A,B)** Examples of expressing FLP-out CsChrimson in smaller numbers of 19f cells. **C)** Percentage of trials that interrupted forward locomotion, stimulated for 6 seconds. The data was separated into two groups where 1-2 19f cells were expressed and where more than 2 19f cells were expressed. **D)** Forward wave frequencies before, during and after 6 second stimulation. **E)** Summary of forward wave durations before, during and after 6 second stimulations. Asterisks indicate significant differences amongst groups (*P < 0.05; one-way repeated measures ANOVA with Bonferroni post-hoc test). Samples sizes: n = 5 (17 trials) where 1-2 19f cells were expressed, n = 6 (15 trials) where more than 2 19f cells were expressed.

## METHODS AND MATERIALS

### Neuron Reconstruction in CATMAID

We used an existing electron microscopy dataset ^63^ to reconstruct larval interneurons and their synaptic partners ^64,65^. Reconstruction of neurons within this dataset was performed using the Collaborative Annotation Toolkit for Massive Amounts of Image Data (CATMAID) ^40^. Neurites of interest were identified and manually traced to build 3D skeletons across serial sections. Synapses were identified by morphological features characteristic of synaptic structures and annotated using ‘connector’ nodes to map synaptic connectivity ^65^.

### Locating Neurons of Interest

Light microscopy images of VT124H03-GAL4-driven GFP expression were generated using a Nikon AZ-C2+ confocal microscope. Samples were fixed, and incubated with primary chicken anti-GFP antibody followed by a secondary Goat anti-chicken 488 antibody. Images were used to locate the approximate position of the neurites of interest within the EM volume.

### Nomenclature and Reconstruction

Neurons were initially named by CATMAID arbitrarily and later annotated based on the GAL4 line or by developmental criteria such as position of cell body and segment. For simplicity, neurons were referred to as ‘19f’ throughout the study. Pre and post-synaptic partners of 19f neurons were reconstructed based on the connectivity data provided by CATMAID, with an emphasis on understanding the connectivity patterns within and across segments.

### Validation and Figure Preparation

Reconstructed neurons were validated against light microscopy data to check for inconsistencies. The CATMAID review widget was used to proofread and adjust reconstructed skeletons. Figures were created using the 3D Viewer and Connectivity widgets in CATMAID and montaged in Inkscape for publication.

### Fly Genetics for Immunostaining

Flies were maintained at 25°C on standard fly food. For experiments requiring Multi-Color-Flp-Out (MCFO) techniques, flies were subjected to a heatshock protocol of 37°C for 15-45 minutes one day before staining. The fly lines used included: UAS-myr-GFP, UAS-MCFO (Nern et al., 2015), UAS-Trans-Tango (Talay et al., 2017), 19f-Split-GAL4 (tsh-AD/CyoTb-RFP ; VT002081.DBD/Tm6B), and 19f-GAL4 (VT124H03-GAL4).

### Staining and Tissue Preparation for Confocal Imaging

Dissections of larval CNSs were performed in physiological saline solution composed of 135 mM NaCl, 5 mM KCl, 2 mM CaCl^2^, 4 mM MgCl^2^, 5 mM TES, and 36 mM sucrose, pH 7.15. CNSs were fixed in 3.7% formaldehyde for 8 minutes at room temperature or on ice when preparing for ChAT staining. Tissues were washed three to four times with phosphate buffered saline (PBS) for 15 minutes each and then incubated in a blocking solution of 1% PBS containing 2% normal donkey serum and 0.1% Triton X-100. Primary antibodies were applied at room temperature for 2 hours, followed by secondary antibodies for 1 hour at room temperature (extended to 48 hours for primary and 12 hours for secondary antibodies at 4°C for ChAT staining). Primary antibodies used were chicken anti-GFP (1:1000, AvesLabs), chicken anti-V5 (1:300, BethylLabs), mouse anti-HA (1:200, BioLegend), and rat anti-FLAG (1:300, NovusBiologicals).

Secondary antibodies were used at a 1:400 dilution from Jackson Immuno Research. Post-staining, tissues were sequentially dehydrated in ethanol solutions, cleared in xylene, and mounted on glass slides with Depex mounting medium. Confocal microscopy was performed using a Zeiss 880 or 800 (Jena, Germany) with 20x or 40x oil objectives, and images were processed using ImageJ and finalized in Inkscape.

### Ca^2+^ Imaging of 19f Cells and aCC Motoneurons

19f-Split-GAL4 flies were crossed with flies expressing the Ca^2+^ indicator GCaMP6f in aCC motoneurons. Isolated 3^rd^ instar larval CNSs were imaged for 10 minutes at 9.94 frames per second with a 100 ms exposure time using a customized BX51 Microscope (Olympus, Tokyo, Japan) equipped with an optoLED light source (Cairn Research Ltd, Faversham, UK) and a iXon EMCCD camera (Oxford Instruments, Belfast, Ireland). Images were acquired with WinFluor imaging software (University of Strathclyde, UK). Regions of interest (ROIs) were adjusted to match the diameters of 19f and aCC cell bodies, with additional ROIs placed on the 19f cell terminal regions to measure synaptic activity. Fluorescence signals from the ROIs were processed using custom scripts in Spike 2 (Cambridge Electronic Design, Cambridge, UK) for signal analysis. Data were plotted using GraphPad Prism 5 and refined for presentation in Inkscape.

### Optogenetic Manipulation of 19f Neurons

The 19f-Split-GAL4 line (exact genotype: tsh-AD/CyoTb-RFP; VT002081.DBD/Tm6B) was utilized in two sets of optogenetic experiments. For activation, the line was crossed to UAS-CsChrimson (UAS-20X-CsChrimson-mvenus in the attP2 landing site). For silencing, it was crossed to UAS-GtACR1 (UAS-GtACR1 in VK00005 landing site). F1 progeny without the Tubby balancer were selected. Experimental larvae were grown in all-trans-retinal supplemented food (0.5 mM, Sigma-Aldrich) until they reached the 3rd instar stage. Control animals were reared on food without retinal supplementation and included animals heterozygous for UAS and GAL4 constructs.

### Electrophysiology

For electrophysiological recordings, larval CNS preparations were dissected in physiological saline solution and pinned in a Sylgard-coated Petri dish. To access the 19f neurons, the ventral nerve cord was exposed using fine dissection tools under a stereomicroscope. A protease solution (0.1-2% Protease XIV, Sigma-Aldrich, St Louis, USA) was applied locally to remove the glial sheath covering the neurons of interest.

Glass electrodes (8-12 MΩ) pulled from borosilicate glass using a P-97 Flaming/Brown micropipette puller (Sutter Instruments, Novato, USA) were filled with a standard internal solution and used for either whole-cell or loose-patch clamp recordings. Recordings were performed using an Axoclamp 2B amplifier (Molecular Devices, San Jose, California, USA), digitized with a Powerlab A-D converter (ADInstruments, Sydney, Australia), and analyzed using LabChart 8.0 software (AD Instruments). Optogenetic stimulation was delivered using a 620 nm LED light source (Cairn Research ltd.), controlled by Chart 7 software to ensure precise timing of light pulses.

### Optogenetic Stimulation Protocol

During electrophysiological experiments, 19f cells expressing either CsChrimson or GtACR1 were optogenetically manipulated using an OptoLED light source (Cairn Instruments) that provided red light pulses (620 nm, intensity ranging between 5-8 mW/mm²). The light was directed to the preparation through a custom-built dichroic mirror housing (Cairn Research Ltd). Red light stimulations were controlled with durations of 6 and 60 seconds using LabChart software.

### Genetic constructs for larval optogenetics

Experimental animals were generated by crossing our 19f-split-GAL4 line to UAS-CsChrimson (20X-CsChrimson-mVenus inserted in the attP2 landing site) for optogenetic activation, and to UAS-GtAcr1 (UAS-GtAcr1 inserted in the VK00005 landing site) for optogenetic inhibition. We also crossed GAL4 and UAS lines to wild-type (Canton-S) animals to generate appropriate heterozygous controls. For optogenetic experiments, animals were maintained in darkness in standard fly food supplemented with 0.5 mM all-trans-retinal (Sigma-Aldrich, Irvine, United Kingdom). Larvae were grown in all-trans-retinal until reaching the 3^rd^ instar stage.

### Behavioral Recording Setup

For behavioral experiments, individual 3^rd^ instar larvae were placed in a square behavior arena (20 cm x 20 cm x 1 cm) layered with a 1% agarose gel (Fisher Scientific, Loughborough, United Kingdom) to facilitate movement tracking. Larval behaviors were recorded using a Ximea mono camera (MQ013MG-E2, Ximea, Münster, Germany), which was integrated into one of the eyepieces of an Olympus dissecting microscope. During recordings, the arena was illuminated with 850 nm infrared (IR) light (OSRAM, Thatcham, United Kingdom) to minimize disturbance to the larvae.

### Optogenetic Stimulation

Freely crawling larvae expressing either CsChrimson or GtACR1, along with relevant controls, were exposed to constant or pulsing (10ms on, 30ms off) red LED light at 617 nm with an intensity of 5-8 mW/mm² (ThorLabs, Exeter, United Kingdom). A 850 nm long-pass filter was used during imaging to maintain consistent image quality during imaging (ThorLabs, Exeter, United Kingdom). Light stimulation and video capture were synchronized and controlled using LabChart software (LabChart 8.0, ADInstruments), which also managed the recording of behavioral responses.

### Behavioral Analysis

Larvae were recorded for periods of 3-4 minutes, with 19f cells stimulated or silenced under constant or pulsing light for durations of 6 and 60 seconds. A minimum interval of 30 seconds was maintained between stimulations to avoid desensitization effects. Behavioral responses were annotated using VCode software ^66^, which allowed for precise timing of various larval behaviors. These behaviors were categorized according to criteria established in previous work ^32^. Data on the durations and frequencies of behaviors were extracted and statistically analyzed using GraphPad Prism software.

### Stochastic Expression of CsChrimson in Subsets of 19f Cells

For stochastic expression studies, UAS-Heatshock-CsChrimson flies (20xUAS-dsFRT-CsChrimson::mVenus, hsFlp2::PEST) were crossed with the 19f-GAL4 driver line. Larvae were raised at 25°C in darkness on food supplemented with 0.5 mM all-trans-retinal (Sigma-Aldrich) and heat-shocked at the 2nd instar stage for 15-30 minutes at 37°C. Post-heat shock, 3rd instar larvae underwent 6-second red light stimulations to activate CsChrimson selectively in subsets of 19f cells. Dissected CNSs were then imaged for mVenus expression to confirm the specific activation patterns.

### Larval Tracking and Olfactory Navigation Experiments

For olfactory navigation assays, 19f-Split-GAL4 flies were crossed to UAS-Kir2.1-GFP flies to achieve neuronal inhibition. A 10 µl droplet of 15 mM ethyl butyrate (Sigma Aldrich, Irvine, United Kingdom) diluted in paraffin oil (Sigma Aldrich) was placed at the center of a 10 cm diameter petri dish layered with a thin agarose gel. Three larvae were placed at the periphery, and their movements toward the odor source were recorded using a Huawei P20 Pro smartphone camera at 60 frames per second. Larval paths were tracked using the TrackMate plugin for Fiji ^67,68^.

### Data Access

All underpinning raw data are freely available upon request. Neuron reconstructions are available in the Virtual Fly Brain website.

## Acknowledgements

We would like to thank Janelia Research Campus for supporting all initial EM reconstruction efforts via the Visiting Scientist Program, and for providing fly lines and constructs. We also thank Cold Spring Harbor Laboratory (CSHL) for space and use of imaging equipment for preliminary experiments as part of the Neurobiology of *Drosophila* Summer course. We also thank Prof. Ellie Hecksher and the Hecksher laboratory for Trans-Tango and MCFO constructs, training, and expert assistance in getting Trans-Tango and MCFO experiments to work. We thank Kevin Myers for assistance with animal tracking software and Prof. Andrew Dacks for providing comments on the manuscript. This work was supported by National Institutes of Health grant (NIH R01-NS105748) awarded to Ellie Heckscher, National Science Foundation Division of Integrative Organismal Systems (NSF IOS 1523125) and National Institute for Drug Abuse (NIDA R13DA034437) grants awarded to Neurobiology of *Drosophila* course at CSHL, by a donation from Kaunas Water Supply (Kaunas, Lithuania), and by a Wellcome Trust Institutional Strategic Support Fund Seed Funding award to SRP.

